# Quantifying community responses to environmental variation from replicate time series

**DOI:** 10.1101/2021.01.12.426425

**Authors:** Joseph S. Phillips, Lucas A. Nell, Jamieson C. Botsch

## Abstract

Time-series data for ecological communities are increasingly available from long-term studies designed to track species responses to environmental change. However, classical multivariate methods for analyzing community composition have limited applicability for time series, as they do not account for temporal autocorrelation in community-member abundances. Furthermore, traditional approaches often obscure the connections between responses at the community level and those for individual taxa, limiting their capacity to infer mechanisms of community change. We show how linear mixed models that account for group-specific temporal autocorrelation and observation error can be used to infer both taxon- and community-level responses to environmental predictors from replicated time-series data. Variation in taxon-specific responses to predictors is modeled using random effects, which can be used to characterize variation in community composition. Moreover, the degree of autocorrelation is estimated separately for each taxon, since this is likely to vary due to differences in their underlying population dynamics. We illustrate the utility of the approach by analyzing the response of a predatory arthropod community to spatiotemporal variation in allochthonous resources in a subarctic landscape. Our results show how mixed models with temporal autocorrelation provide a unified approach to characterizing taxon- and community-level responses to environmental variation through time.

## Introduction

A central goal of ecology is to understand how environmental variation affects population and community dynamics. To this end, time-series data of ecological communities are increasingly available from monitoring programs such as the National Ecological Observatory Network (Keller et al. 2008) and long-term field experiments (e.g., Fraser et al. 2013, Cowles et al. 2018). Data of this sort provide great promise for answering long-standing questions of how environmental factors drive community dynamics. The responses of ecological communities to environmental predictors are often analyzed using ordination methods (e.g., NMDS, RDA), which map variation in abundance or occurrence onto orthogonal axes that provide synoptic assessments of community variation (McGarigal et al. 2013). However, when applied to community time series these approaches do not account for temporal autocorrelation in the abundances of the populations that compose the community. This is an important limitation, as failing to account for temporal autocorrelation can lead to erroneous conclusions (Ives and Zhu 2006). Furthermore, populations may vary in their degree of temporal autocorrelation, due to differences in life histories or strengths of population regulation (Ziebarth et al. 2010), potentially exacerbating the challenges for statistical inference. One approach to this problem is to use dimension reduction to produce composite community metrics for more traditional univariate time-series analyses (Simpson and Anderson 2009). Unfortunately, this approach sacrifices valuable inferences about taxon-specific responses to environmental predictors. Alternative approaches are designed to explicitly quantify dynamic interspecific interactions but require relatively complex models and long time series (Ives et al. 2003, Hampton et al. 2013). In contrast, community time series are often relatively short but contain replication through space or across experimental units with the goal of inferring the responses of communities to external drivers or experimental manipulations.

Here, we show how autoregressive mixed models (linear mixed models with temporal autocorrelation structures) that account for observation error can be used to infer taxon- and community-level responses to predictor variables from replicated time series data. The approach is based on the method of Jackson et al. (2012), who model variation in taxon-specific responses to predictors as random effects. These random effects can in turn be used for statistical inferences on community composition, which fundamentally derives from variation among taxa, making explicit the link between the two organizational scales. We extend this approach for time series by formulating models with separate temporal autocorrelation structures for different taxonomic groups, accounting for the fact that different taxa are likely to be characterized by different dynamics. The degree of temporal autocorrelation itself provides information about the community dynamics (Ives et al. 2003), and its inclusion allows for valid statistical inferences on the effects of predictors when observations are temporally replicated (Ives and Zhu 2006).

We illustrate the utility of this method by analyzing the responses of terrestrial arthropods to spatiotemporal variation in allochthonous resources at Lake Mývatn, Iceland (Einarsson et al. 2004). Mývatn has large populations of midges (Diptera: Chironomidae) that emerge from the lake as adults and subsequently subsidize the terrestrial plant (Gratton et al. 2008, Krowiak et al. 2017) and arthropod (Dreyer et al. 2012, Sanchez-Ruiz et al. 2018) communities. The midges have large interannual fluctuations in abundance and decline in deposition with distance from the lakeshore (Dreyer et al. 2015). We apply autoregressive mixed models to assess the community response to the highly variable midge subsidy, while simultaneously accounting for linear trends across time and distance from the lake. Including those two covariates was important for isolating the effects of midges from those of other variables that vary with distance (e.g., plant composition) or time (e.g., climate) (Bowden et al. 2018). This study shows how autoregressive mixed models provide a statistically sound, easy-to-interpret method for characterizing community responses to environmental variation in time and space.

## Methods

Mývatn has a tundra-subarctic climate, and the surrounding landscape is dominated by heathland and grassland vegetation (Einarsson et al. 2004). Arthropod samples were collected approximately every 15d from five sites around the shoreline of the lake from May–August in 2008–2019. Each site included 3–4 plots at various distances (5m, 50m, 150m, and 500m) along a transect perpendicular to the lakeshore (some transects only extended to 150m). Midge deposition was sampled at each site using aerial infall traps (Dreyer et al. 2015), and activity-density of ground-dwelling arthropods was sampled using pitfall traps (Southwood and Henderson 2009).

Activity-density is a measurement of relative abundance, which represents a combination of both population size and behavioral phenomena, both of which potentially respond to the allochthonous inputs of resources (Ostfeld and Keesing 2000). For this analysis, we used the cumulative May–August catch of predatory and detritivorous arthropods (Figure 1), including ground beetles (Carabidae), rove beetles (Staphylinidae), harvestmen (Opiliones), ground spiders (Gnaphosidae), sheet-weaving spiders (Linyphiidae), and wolf spiders (Lycosidae).

**Figure 1.**
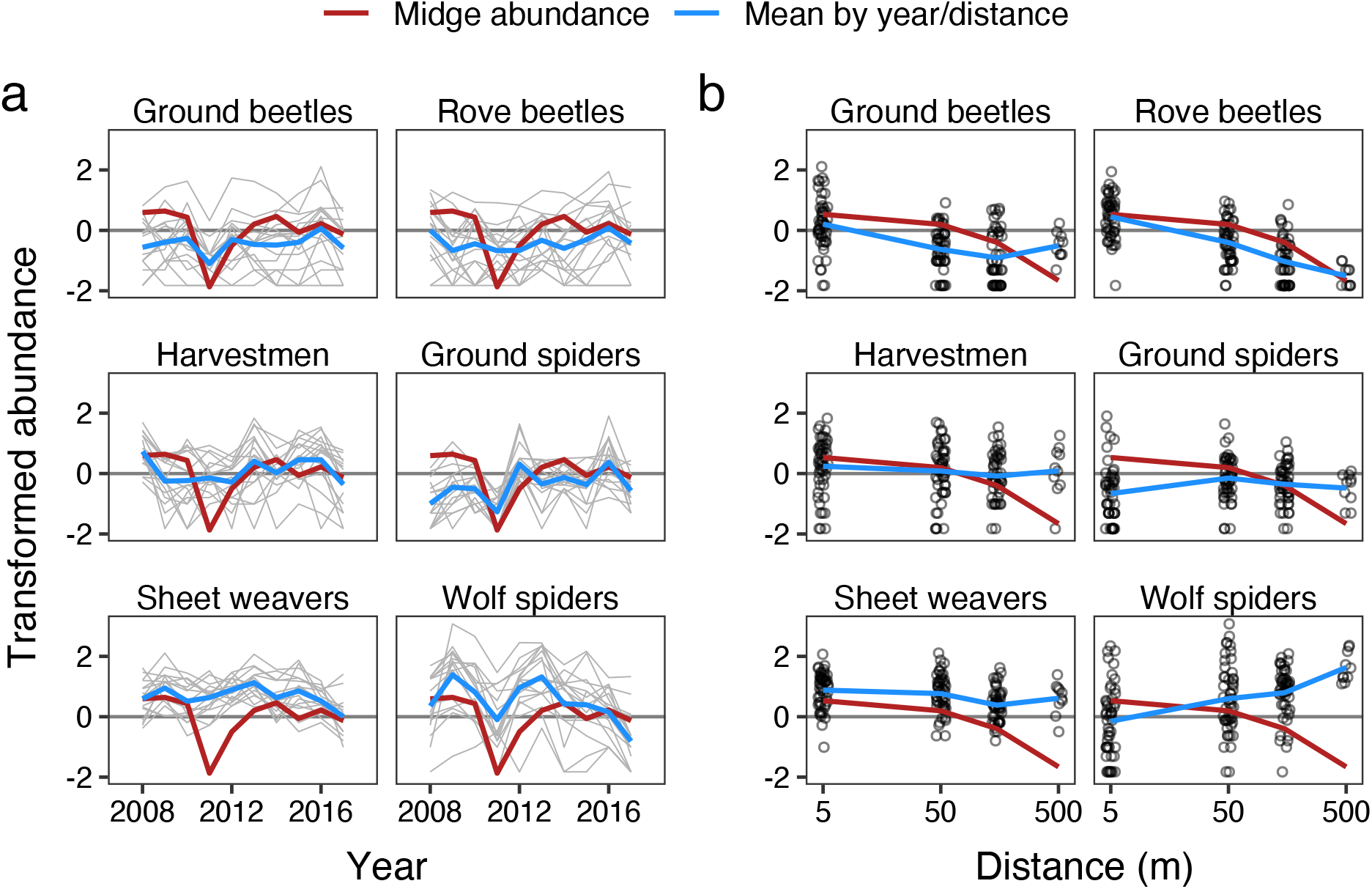
Community time series data for 6 predatory arthropod taxa at Lake Mývatn. Transformed abundance of arthropods (a) through time and (b) by distance from the lake. (a) Narrow gray lines are time series grouped by site and distance. Thick red lines are transformed midge abundances averaged by (a) year or (b) distance. Thick blue lines are transformed abundances averaged by taxon and (a) year or (b) distance. Relative abundances were measured using activity-densities that were log-transformed, then z-scored across the entire data set.

We analyzed taxon-level variation using an extension of linear mixed models. Our model included (1) fixed effects for continuous predictor variables (number of years since initial sampling year, distance from the lake, and midge deposition); (2) random intercepts grouped by taxon, taxon × site, and taxon × plot; and (3) random slopes for the three predictor variables grouped by taxon. The fixed effects give the average response across taxa to the predictor variables, while the corresponding random effects give the deviation for each taxon from this average response (Jackson et al. 2012). The overall response of a given taxon is the sum of the fixed and random components. Quantifying responses for each taxon in a single model allows for “shrinkage” or “partial pooling” of the taxon-specific responses towards the mean response, which reduces the noisiness of individual estimates and ameliorates concerns of multiple comparisons (Gelman et al. 2012) that have been raised for the examination of responses for many taxa separately (McGarigal et al. 2013). We log-transformed midge deposition and distance before z-scoring all of the predictors by either subtracting the mean (midges, distance) or the minimum (time) and then dividing by the standard deviation. For time, we subtracted the minimum so that the intercepts quantified the initial abundance of each taxon in each plot.

To formulate the autoregressive component of the model, we defined a vector **y** consisting of transformed counts for each taxon in each plot through time. Because population processes are generally multiplicative, we log-transformed the counts, adding one to all values to accommodate the modest number of zeros. We then z-scored all values to improve interpretability of the model coefficients. The rows of **y** were grouped as individual time series (i.e., taxon–plot combinations), with observations within each group ordered by time. For each time series, we then specified an autoregressive state-space model for the *i*th observation of **y** as

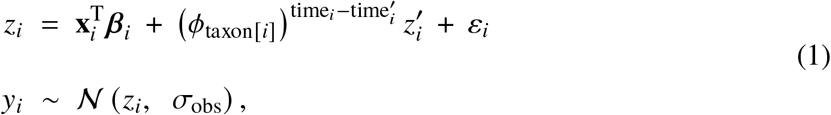

where *z*_*i*_ and 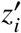 are latent and lagged-latent abundances on the transformed scale, 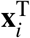 is the transposed vector of predictor values for the *i*th observation, *β*_*i*_ is a vector of coefficients modeled hierarchically according to the fixed and random effects structure described above, *ϕ*_taxon[*i*]_ is the taxon-specific autoregressive (AR) parameter, time_*i*_ is the current time (in years), 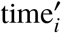 is the lagged time, *ε*_*i*_ is the Gaussian process error with standard deviation *σ*_proc_, and *σ*_obs_ is the observation error standard deviation. Because *y*_*i*_ consists of log-transformed abundances, equation 1 is essentially a log-linear model of population dynamics, with the population growth rate given by 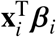. When *ϕ*_taxon[*i*]_ = 0, the dynamics are determined primarily by 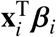, indicating strong population regulation associated with contemporary environmental conditions (Ives et al. 2010, Ziebarth et al. 2010). In contrast, *ϕ*_taxon[*i*]_ near 1 implies weak regulation as there is strong “memory” of previous population states (i.e. high temporal autocorrelation). Exponentiating the elapsed time with base *ϕ*_taxon[*i*]_ allows the model to accommodate unequal time steps (Zuur et al. 2009). The autoregressive term in equation (1) was dropped for the initial value of each time series, such that the initial value was given entirely by predictor variables and associated coefficients. The primary difference between the present approach and more traditional linear models with AR structures (e.g., Zuur et al. 2009) is the estimation of separate values of the AR parameter for different groups (i.e., taxa). This is an important extension, as it allows members of a community with different levels of temporal autocorrelation to be included in a single model.

Community analyses often utilize dimension reduction to characterize variation in community composition (McGarigal et al. 2013). However, with autoregressive mixed models it is possible to infer variation in community composition directly from the variation in taxon-specific responses. We illustrated this in two ways. First, we examined the posterior distributions of the fixed and random effects associated with the time, distance, and midge deposition. Fixed effects characterize the mean responses among taxa, and substantial overlap with zero would indicate the absence of a consistent community-wide response to a given predictor. Random effects characterize variation in the responses among taxa, which is the basis for changes in composition per se. While random effect standard deviations cannot overlap zero, concentration of posterior density near the zero-boundary would imply limited variation in community composition in response to a given predictor. Second, we used principal components analysis on fitted values from the model to generate orthogonal axes characterizing the expected variation in the community due to the predictors (similar to Jackson et al. 2012), and we used ANOVA to partition variation in the fitted values into contributions from midges, time, and distance. We then projected the observed data onto the PC axes. The total contribution of each predictor to the observed community variation was calculated as the sum of the axis loadings weighted by the corresponding variance contributions. These two approaches provided statistical inference and visualization of community abundance and composition analogous to conventional ordination while appropriately accounting for temporal autocorrelation and maintaining an explicit link to taxon-specific responses.

We fit the model with a Bayesian approach using Stan, via the rstan (Stan Development Team 2020) package in R 4.0.3 (R Core Team 2020). We used 4 chains with 4000 iterations each (including 2000 iterations of “warm-up”) and assessed convergence based on effective sample size across the Markov chains, number of divergent transitions, and potential scale reduction factor. We used Gaussian priors with mean 0 and standard deviation 1 for the fixed effects, Gamma priors with shape 1.5 and rate 3 for the standard deviations, and a Gaussian prior with mean 0 and standard deviation 0.5 for the AR parameters. The latter was truncated with a lower bound of zero to avoid quasi-cyclic dynamics, but did not have an explicit upper bound. We used posterior medians as point estimates and 68% quantiles as bounds of uncertainty intervals, matching the coverage of standard errors (hereafter “posterior standard errors”). Data and code can be found at https://github.com/lucasnell/trans_trends and https://github.com/lucasnell/trans_trends_pkg.

## Results

The AR coefficients were generally similar across taxa and all well below 1 (Figure 2a), indicating that the populations were statistically stationary through time after accounting for the predictors (including the linear time trends). Ground spiders had the lowest AR coefficient; this was visually apparent in the data, as ground spiders appeared more tightly constrained to their mean abundance than was the case for other taxa (Figure 1a). This implies that ground spiders may have experienced the strongest population regulation as determined by the environmental predictors, although caution is warranted when inferring dynamical features of short time series.

**Figure 2.**
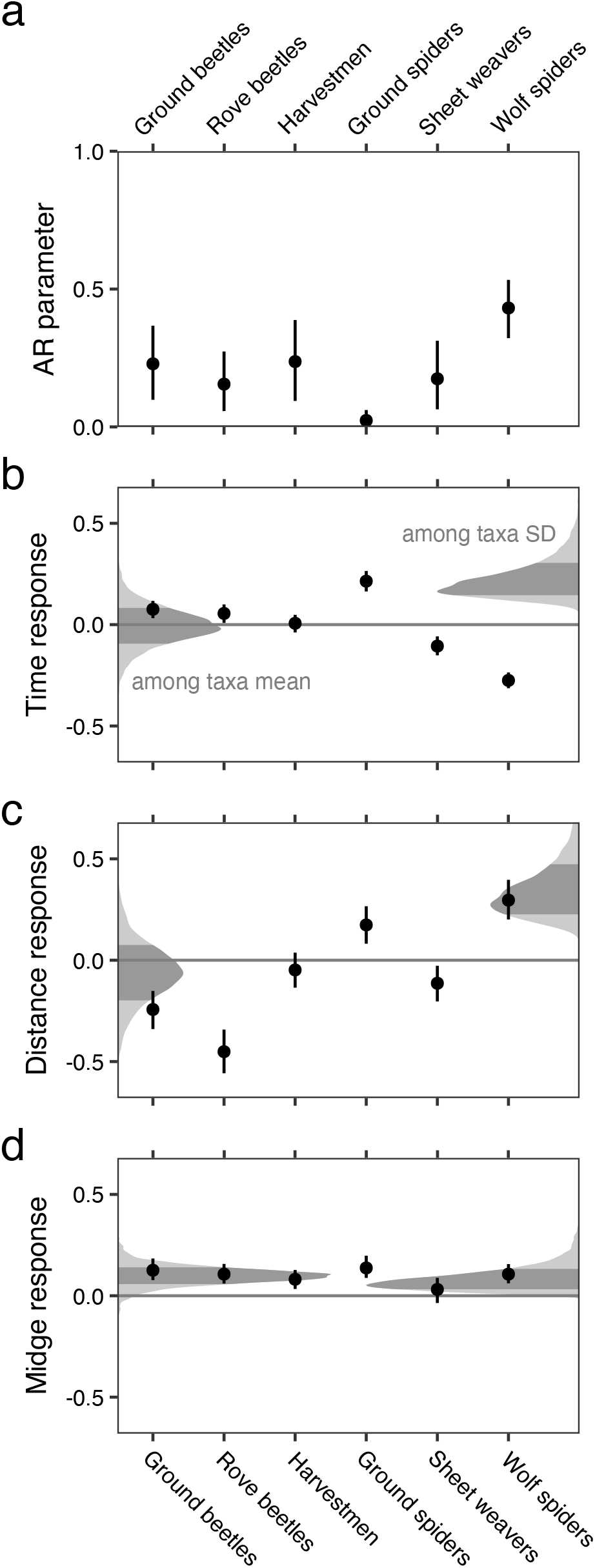
Parameter estimates from the autoregressive mixed model, including (a) autoregressive coefficients for each taxon, and (b–d) responses to each predictor. In all panels, points are posterior medians, error bars are posterior standard errors. (b–d) The shaded curves on the left show the posterior densities for the mean response across taxa (i.e., fixed effects), and shaded curves on the right show posterior distributions for the random effect standard deviations; darker shaded regions indicate posterior standard errors.

Taxa varied substantially in their trends through time: ground spiders and beetles increased, wolf spiders and sheet weavers decreased, while rove beetles and harvestmen remained largely unchanged (Figure 2b). Variation among taxa was even greater in response to distance than for time (Figure 2c), with ground and rove beetles most abundant near the lake, while ground and wolf spiders were most abundant further away. In contrast to time and distance, the average response across taxa to midge deposition was consistently positive (although fairly weak). Indeed, the responses for all individual taxa were positive, with only sheet weavers approaching zero (Figure 2d). The positive responses to midges were associated with variation in deposition both among years and along distance from the lake (Figure 1), underscoring the importance of accounting for time and distance when inferring responses to midge deposition.

The taxon-specific responses provided the basis for inferring variation in both community composition per se and community-wide abundance. Specifically, changes in composition arose when taxa differed in their responses to predictors, while changes in community-wide abundance arose from consistent responses among taxa. For example, taxa differed in their responses to distance from the lake (“among taxa SD” in Figure 2c), which in turn implied that community composition varied with distance. In the PCA, this resulted in substantial variation among the taxon-response vectors along PC1, which was most strongly associated with distance (Figure 3a–d,g). Taxa also differed in responses to time (“among taxa SD” in Figure 2b), with variation in community composition manifesting along PC3 (Figure 3c–f,g). In contrast, the average response to midges was positive (“among taxa mean” in Figure 2d), resulting in an increase in community-wide abundance along PC2 (Figure 3a,b,e,f,g). Distance accounted for most of the observed community variation, followed by midges, and finally time (Figure 3g). Together, linear responses of the various taxa to midges, distance, and time accounted for 74% of the observed community variation when projected onto the PC axes.

**Figure 3.**
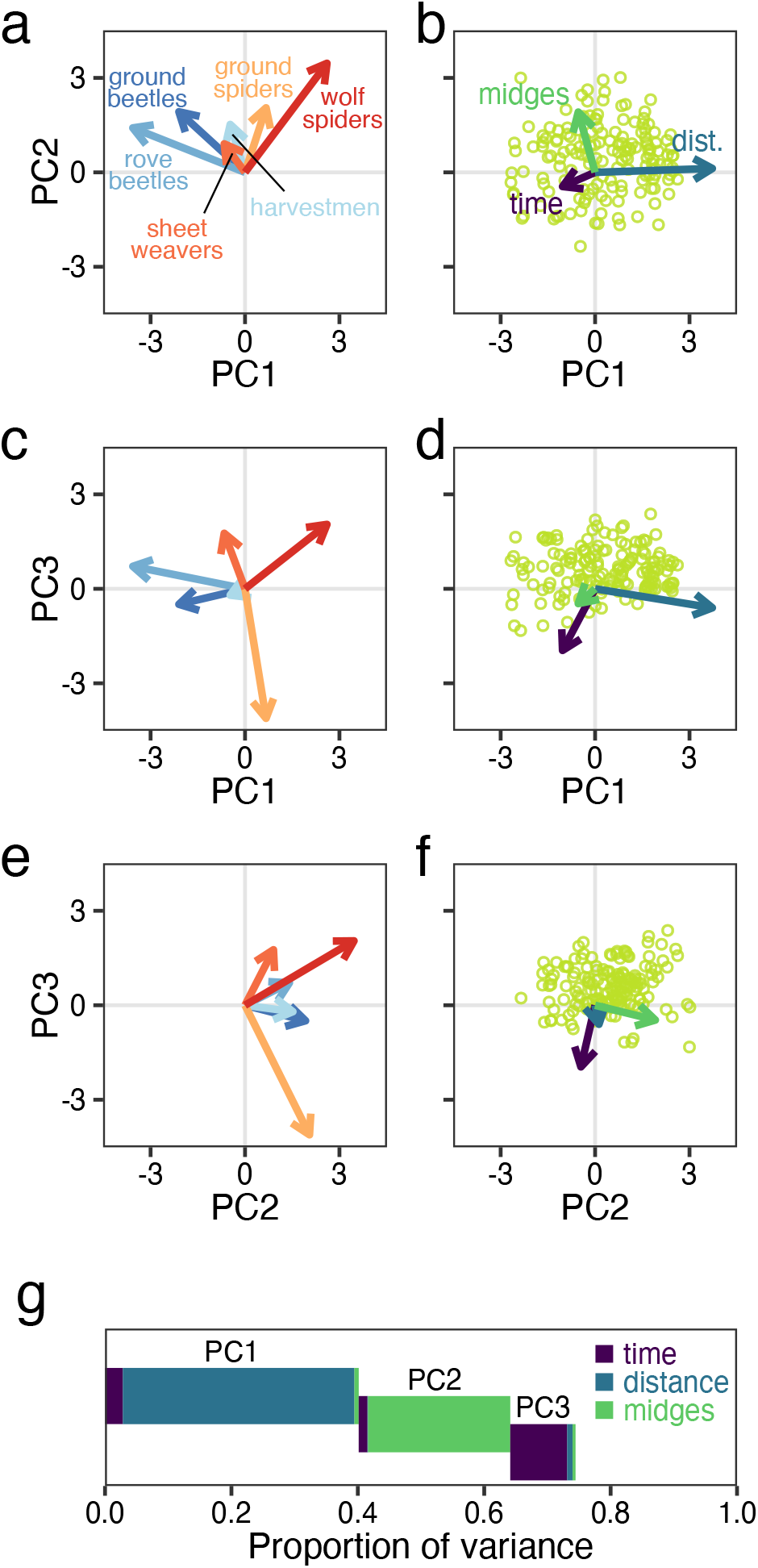
Principal components analysis (PCA; a–f) and variance partitioning (g). The PCA is based on the taxon-specific responses to time, distance, and midges, as inferred from the model. The bi-plots show (a,c,e) taxon-response vectors and (b,d,f) model predictors and observed data projected onto the PC axes. (a–f) Panel rows separate the three pairwise combinations of PC axes. (b,d,f) Points indicate observed data. For clarity of visualization, the taxon vector overlays in panels a,c,e are scaled relative to vectors in b,d,f, and both are scaled relative to observed data projections. (g) Variance partitioning of the observed data projected onto the PC axes. Each PC axis was partitioning into contributions from the model predictors (time, distance, and midge deposition) through ANOVA. Portions of the variance in the observed data explained by a given PC axis are offset vertically, while the the variance in the observed data explained by a given predictor are indicated by color.

## Discussion

Here, we show how linear mixed models with temporal autocorrelation can be used to quantify community responses to environmental variation from replicated time series. The approach extends the previous use of mixed models for estimating taxon- and community-level responses to environmental variation (Jackson et al. 2012, Bartrons et al. 2015) by incorporating temporal autocorrelation with group-specific values for the autoregressive parameter. While our model formulation accounted for space using linear trends (due to the nature of the example data), the method could be extended to explicitly include spatial autocorrelation when appropriate (similar to Bartrons et al. 2015). The approach of modeling community responses at the taxon level provides a natural way to accommodate the temporal autocorrelation structure of community times-series data, because community dynamics are manifestations of the dynamics of constituent populations. Furthermore, the taxon-specific autoregressive parameter estimated by our approach has a clear population-dynamic interpretation as the strength with which the population is drawn towards its central tendency, which is closely tied to classical ecological concepts of population regulation (Nicholson 1933, Ziebarth et al. 2010). In our application to the Mývatn arthropod community, the AR parameters for all taxa were well below 1, indicating that the populations were statistically stationary through time after accounting for linear time trends and responses to midge deposition. Taxa also varied substantially in their AR parameters, with the dynamics of ground spiders appearing to be more tightly constrained than for the other taxa. However, caution is warranted when drawing such inferences from short time series and when not explicitly accounting for interspecific interactions.

While our approach is formulated in terms of taxon-specific responses, we illustrate two methods for how these can be extended to draw inferences about community composition. First, the model provides estimates of both the mean response (fixed effects) and the variation in responses (random effects) among taxa. When posterior distributions for the means or standard deviations are concentrated away from zero, this indicates substantial variation in community-wide abundance and composition per se, respectively. Second, principal components analysis can be applied to the fitted values from the model (similar to Jackson et al. 2012), which allows variation in community composition to be quantified in a manner analogous to traditional ordination while appropriately accounting for temporal autocorrelation. The variance in the principal components analyses can then be partitioned into contributions from the predictor variables. This approach exemplifies the general dictum that it is better to ‘analyze, then aggregate’ than it is to ‘analyze the aggregate’ (Clark et al. 2011). In our example application to the Mývatn data, most of the community variation was due to changes in composition per se resulting from heterogeneous taxon-specific responses to distance from the lake and across time. However, most taxa were positively associated with midge deposition, resulting in elevated abundance across the entire predatory arthropod community.

This study fits into a larger effort to understand the role of allochthonous inputs on recipient communities (Polis et al. 1997, McCary et al. 2020). One of the challenges in identifying the effects of allochthonous pulses from long-term observational data is that pulses are likely to be conflated with other factors that vary through space and time (Yang et al. 2008). By using linear mixed effects models with group-specific temporal autocorrelation, we were able to partition the influence of three interrelated variables (time, distance, and midge deposition) to quantify the effect of allochthonous resources on a tundra arthropod community. We found a positive overall response of predator and detritivore abundance to midge deposition, which is consistent with previous studies at Mývatn (Hoekman et al. 2011, Dreyer et al. 2012, Sanchez-Ruiz et al. 2018) and in other systems (Ostfeld and Keesing 2000, Murphy et al. 2012). Given the ubiquity of spatiotemporal associations between different drivers of ecological communities, our approach has potential utility across many subdisciplines of community ecology.

## Acknowledgments

Data collection was supported by National Science Foundation grants DEB-0717148 to Claudio Gratton, DEB-1052160 and −1556208 to Anthony R. Ives, and Graduate Research Fellowships DGE-1256259 and −1747503. We thank Árni Einarsson and Mývatn Research Station for research support, and Amanda McCormick, Matthew McCary, Claudio Gratton, and Anthony R. Ives for feedback on the manuscript. LAN and JSP made equal contributions to this work.

